# Statistics to prioritize rare variants in family-based sequencing studies with disease subtypes

**DOI:** 10.1101/2023.09.28.560053

**Authors:** Christina Nieuwoudt, Fabiha Binte Farooq, Angela Brooks-Wilson, Alexandre Bureau, Jinko Graham

## Abstract

Family-based sequencing studies are increasingly used to find rare genetic variants of high risk for disease traits with familial clustering. In some studies, families with multiple disease subtypes are collected and the exomes of affected relatives are sequenced for shared rare variants. Since different families can harbor different causal variants and each family harbors many rare variants, tests to detect causal variants can have low power in this study design. Our goal is rather to prioritize shared variants for further investigation by, e.g., pathway analyses or functional studies. The transmission-disequilibrium test prioritizes variants based on departures from Mendelian transmission in parent-child trios. Extending this idea to families, we propose methods to prioritize rare variants shared in affected relatives with two disease subtypes, with one subtype more heritable than the other. Global approaches condition on a variant being observed in the study and assume a known probability of carrying a causal variant. In contrast, local approaches condition on a variant being observed in specific families to eliminate the carrier probability. Our simulation results indicate that global approaches are robust to mis-specification of the carrier probability and prioritize more effectively than local approaches even when the carrier probability is misspecified.

## 1 Introduction

Family-based sequencing studies are employed to identify causal rare variants, or *c*RVs, for a disease. They can involve a handful of selected families that are highly enriched for cases or several dozens of families with more than one case obtained from a population-based disease registry. In these studies, genetic sharing among disease-affected relatives with shared traits, such as disease severity levels or disease subtypes, is of interest. In this work, we consider diseases with two distinct subtypes, or with more than two subtypes that can be classified into two distinct groups, where one group of subtypes is more heritable than the other based on expert advice or existing diagnostics (e.g. subtype-specific recurrence risks or heritability estimates). For example, among lymphoid cancers, Hodgkin Lymphoma (HL) is considered to be a more genetic subtype than Non-Hodgkin Lymphoma (NHL) since first degree relatives of NHL-affected relatives experience a 1.8-fold elevated risk of developing NHL, whereas first degree relatives of HL-affected relatives experience a 3.3-fold elevated risk of developing HL [Cerhan and Slager, 2015].

The methodology presented in this work is developed primarily to address the needs of family-based studies with characteristics similar to the Lymphoid Cancer Families Study at BC Cancer. Consenting families are ascertained for multiple relatives affected by lymphoid cancer; that is, Hodgkin lymphoma, non-Hodgkin lymphoma, lymphocytic leukaemia, or myeloma. Then the exomes of the disease-affected relatives are sequenced. When available, informative unaffected relatives, such as unaffected and presumably non-transmitting parents will be included to rule out a subset of variants. Occasionally, suspected obligate carriers may be sequenced to resolve questions about variant frequency or transmission of the variants among relatives. However, most sequenced subjects in the study are expected to be disease-affected relatives. We therefore assume, throughout, that only disease-affected relatives are sequenced. Additionally, since the majority of families contain affected relatives separated by very few meioses we anticipate limited statistical power to detect causal variants. Furthermore, we suspect extensive locus heterogeneity of lymphoid cancer [e.g., Cerhan and Slager, 2015]. As a result, we do not expect the same causal variants, or even different causal variants within the same gene, to be shared between different families. Since genetic linkage or identity-by-descent sharing approaches are not well-suited to locus heterogeneity [Ott et al., 2015] we look to alternative approaches to prioritize variants for follow-up. In this work, we aim to define variant-specific sharing statistics that account for kinship amongst the affected family members and observed subtype of disease. These variant-specific statistics may be used as ranks or scores in future analyses of pathway enrichment or more labor-intensive functional studies. To develop variant-specific sharing statistics, we will build on existing statistical methods.

Basu et al. [2010] construct a likelihood-based test to detect linkage between a genetic marker and a disease of interest. They introduce a preferential transmission model to test the null hypothesis of no linkage between the disease and the genetic marker against the alternative hypothesis that the disease and the marker are linked. Their preferential transmission model is parametrized by trait-specific transmission probabilities and so is able to use information provided by unaffected relatives. Since the authors consider binary traits (i.e. affected/unaffected) two transmission probabilities are introduced: one for affected individuals and one for unaffected individuals. The preferential transmission model is similar to the gamete-competition model [Sinsheimer et al., 2000] in that, through the transmission probabilities, the alleles of the parent compete for transmission to the offspring. However, the gamete-competition model aims to identify which alleles are associated with a particular trait whereas the preferential transmission model aims to identify which loci are linked to a trait. The preferential-transmission and gamete-competition models are more parsimonious than traditional parametric linkage or general allelic transmission models, respectively. Parsimony is important because of the potential for over-parametrized models leading to unstable inference and ambiguous interpretation, and because the computational burden increases substantially with the number of parameters. Bureau and colleagues (Bureau et al. [2014a,b, 2019]) present exact-probability methodology to test for excess sharing of rare variants observed in disease-affected relatives. They consider variants that are shared between all sequenced affected relatives in a pedigree assuming that the variant is rare enough to have been introduced by at most one founder. Bureau et al. [2019] later extended this work to allow for sharing among a subset of the disease-affected relatives in a pedigree.

Motivated by the transmission models of Sinsheimer et al. [2000] and Basu et al. [2010] and the rare-variant sharing (RVS) methods of Bureau (Bureau et al. [2014a,b, 2019]), we propose a likelihood-based approach to prioritizing rare variants shared by affected family members. Following Bureau and colleagues, we focus on the genetic information of the disease-affected relatives and extend their methodology to allow for trait-specific transmission parameters as in Basu et al. [2010] and Sinsheimer et al. [2000]. Our alternative model is parameterized by subtype-specific transmission probabilities of rare variants. The model leads to several statistics to rank rare variants observed among the disease-affected relatives in a study. These statistics take high values when excess sharing is observed, especially when the sharing occurs between relatives affected with a disease subtype that is more heritable. Additionally, these statistics increase as the kinship or degree of relatedness decreases between disease-affected relatives in which sharing has been observed. To our knowledge, the statistics we propose are the first to incorporate the information from disease subtypes into the analysis of rare variants in affected-only family designs.

The remainder of the article is structured as follows. In Section 2 we develop statistics for ranking and testing rare variants, and describe the design of a simulation study to compare the methods. In Section 3 we compare the methods in an example data analysis of a simulated dataset. We also present the results of the simulation study. In the example data analysis, we compare the methods through their rankings of the rare variants and their p-values for the corresponding test of association, with a focus on the causal rare variants. For the simulation study, we compare the methods through their median normalized ranks over the simulation replicates, their estimated Type-I error rates and their estimated powers. We conclude in Section 4 with a discussion and directions for future work.

## 2 Models and Methods

Our goal is to develop likelihoods for subtype-specific transmission parameters using data on individual rare variants (RVs) observed in sequenced family members affected with a disease of interest. These likelihoods lead to statistics for ranking and testing rare variants. The model underlying the likelihoods considers the collection of causal RVs as a group. We assume that the group of causal RVs (cRVs) is rare enough that at most one copy of one cRV is introduced into the family through one founder. Consequently, there is at most one copy of any cRV in each affected individual. Likelihoods are thus based on the sharing configurations of individual RVs.

### 2.1 Model

The data are the observed sharing configurations in the pedigrees; that is, the rare variant statuses of the sequenced affected relatives. Throughout, we condition on the number of pedigrees in the study, their family structure and the disease-affection status of the relatives.

The rare-variant status of the sequenced affected individuals is taken to be random. Assume that a study contains *m* unrelated pedigrees and let *n*_*i*_ denote the number of relatives who are affected by either disease-subtype *A* or disease-subtype *B* in pedigree *i*, for *i* ∈ {1, 2, …, *m*}. Without loss of generality, assume that, *a priori*, disease subtype *A* is more genetic than disease subtype *B*. In this work we do not consider sequence data from relatives currently unaffected by disease or suspected obligate carriers. Every sequenced individual is assumed to be affected by one of two disease-subtypes.

For pedigree *i* of the study, the **familial configuration** is denoted 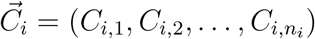, where *C*_*i*,*j*_ is a random variable that denotes the number of autosomal rare variant (RV) copies observed in affected member *j* of pedigree *i*. For example, in the simulated pedigree of Figure 1, ascertained for lymphoid cancer (HL, NHL), suppose that an RV is present in the sequenced affected members with IDs 38 and 49 but not in those with IDs 17 and 29. When the elements of the configuration vector are ordered by the ID numbers of the affected relatives (i.e. 17, 29, 38 and 49) the familial configuration is 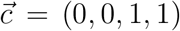. We use the term **global configuration**,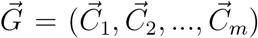, to describe RV configurations spanning every pedigree in the study. We obtain the global configuration by concatenating the familial configurations in the order of their family ID numbers. While a particular rare variant may not be observed in every pedigree, each rare variant under consideration must have been observed in at least one disease-affected subject in the study; i.e. 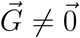.

**Figure 1:**
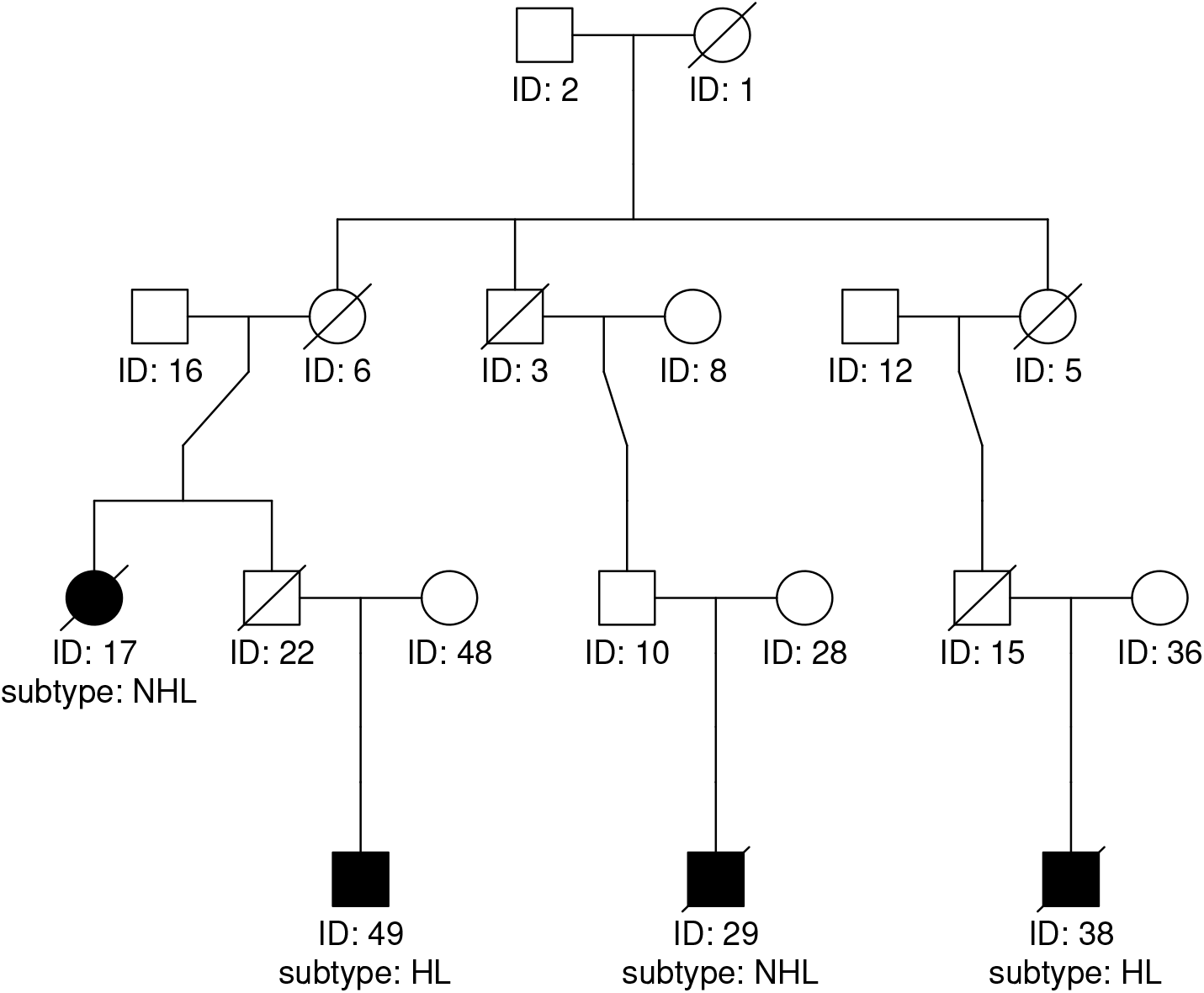
Simulated example family

Now consider a probability model for the global configurations. When families are unrelated, we assume the familial configuration in family *i* is independent of the familial configuration in family *j* and write the probability of a global configuration as a product of the probabilities of the component familial configurations.

To obtain the probabilities of the familial configurations, define the carrier probability, *p*_*c*_, to be the probability of all RVs of interest considered as a group; i.e., causal RVs (*c*RVs) in a gene, genomic region or biological pathway [Nieuwoudt et al., 2018]. Let *n*_*f*,*i*_ denote the number of founders in pedigree *i* and *H*_*i*_ denote the number of *c*RV copies that are introduced to pedigree *i*. Assuming the pedigree founders are independent, *H*_*i*_ has a binomial distribution with *n*_*f*,*i*_ trials and success probability *p*_*c*_. We assume that *p*_*c*_ is small enough that *P*(*H*_*i*_ *>* 1) is negligible, 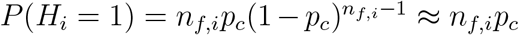 and *P*(*H*_*i*_ = 0) ≈ 1 − *n*_*f*,*i*_*p*_*c*_.

The probability of the familial configuration is then:

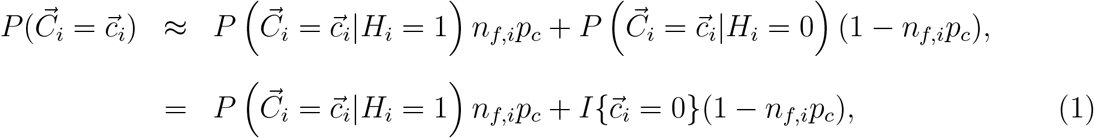

where 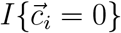 is equal to 1 if 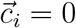 and 0 otherwise. The probability 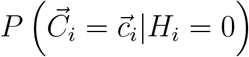 in the second term of the right-hand side reduces to 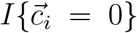 since the only possible configuration when no RV is introduced into the family is 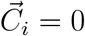.

To derive an expression for the other probability 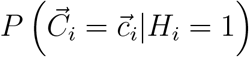 in the right-hand side of equation (1), we define *F*_*i*,*l*_ to be equal to 1 if founder *l* from pedigree *i* is a carrier of the RV and 0 otherwise. Since *P*(*H*_*i*_ *>* 1) ≈ 0 we assume no two *F*_*i*,*l*_ may simultaneously be 1. We further assume that each founder is equally likely to introduce a *c*RV, so that

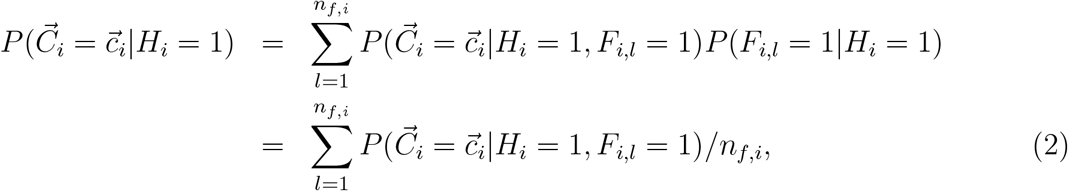

The conditional configuration probabilities 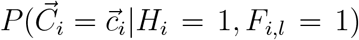 in equation (2) may be parametrized by transmission probabilities. Let *τ*_*i*,*j*_ be the probability that a parent who is heterozygous for a *c*RV transmits it to subject *j* of pedigree *i*. We assume that the transmission probability is disease-subtype dependent so that

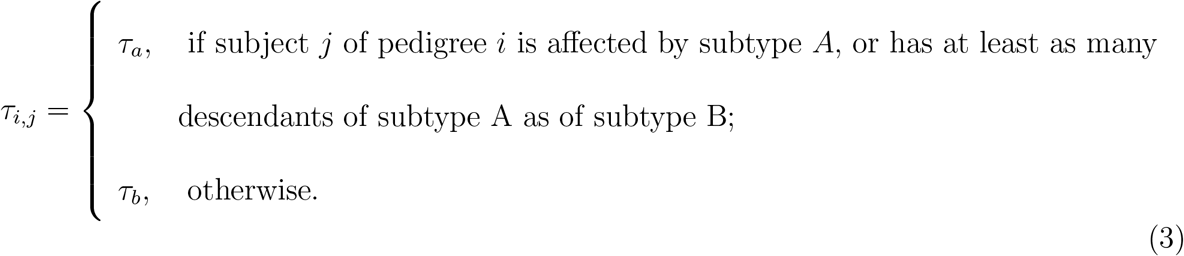

To clarify the relationship between the transmission parameters *τ*_*i*,*j*_ and 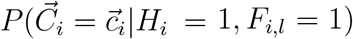 in equation (2), we view the pedigree of affected individuals and their ancestors as a Bayesian network [Lauritzen and Spiegelhalter, 1988, Højsgaard, 2012]. In this context, the Bayesian network is a graph of nodes and edges that represents dependence between pedigree members. Nodes correspond to individuals, edges are defined by transmissions from parents to their offspring and the conditional probability distributions of nodes are determined by the transmission parameters *τ*_*a*_ and *τ*_*b*_ described in equation (3). In the Bayesian network, 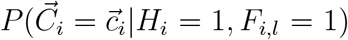 is the conditional probability of the nodes of the sequenced affected individuals given the *c*RV status of the nodes of founders. We write 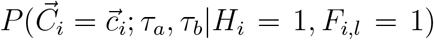 for this probability to emphasize that it depends on the transmission parameters but not the carrier probability *p*_*c*_. Though there is no closed-form expression, 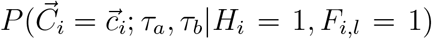 can be calculated for any (*τ*_*a*_, *τ*_*b*_) by software for Bayesian networks, such as the R package `gRain` [Højsgaard, 2012].

To summarize, we can parametrize the familial configuration probabilities by the transmission parameters (*τ*_*a*_, *τ*_*b*_) and the carrier probability *p*_*c*_. In what follows we assume that the carrier probability is known and suppress it in the notation. Thus, the configuration probabilities are denoted 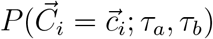. Combining equations (1) and (2) and simplifying yields

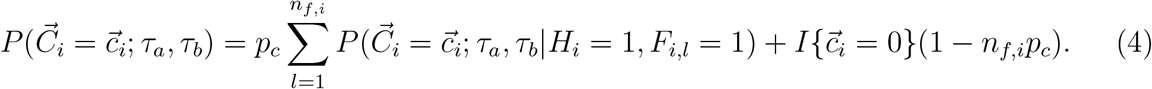

For future reference, equation (4) implies that

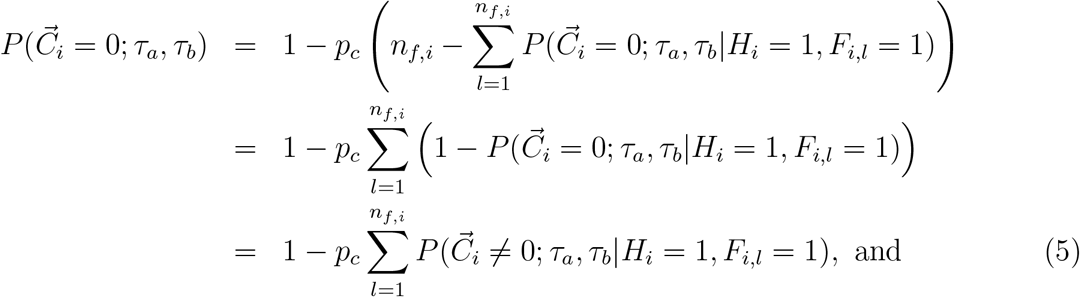

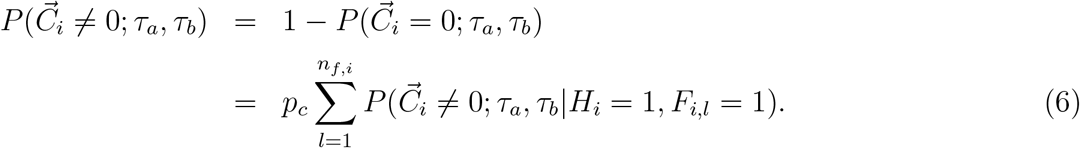

The transmission parameters *τ*_*a*_ and *τ*_*b*_ relate to our null and alternative hypotheses about linkage and association. Following others [e.g., Bureau et al., 2014a, Laird and Lange, 2008], we adopt a null hypothesis of no linkage and no association between the observed variant and the disease-susceptibility locus. Since family-based studies have no power to detect association without linkage [Laird and Lange, 2008], we also assume that under the alternative hypothesis both linkage and association are present.

Under the null hypothesis of no linkage and no association, the disease subtypes are independent of the global configuration, 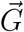, and the parameter space is 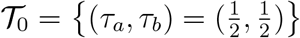. Basu et al. [2010] develop a pedigree-linkage method that considers transmission probabilities to disease-affected and healthy individuals, which we denote by *τ*_*d*_ and *τ*_*h*_, respectively.

Their alternative hypothesis of linkage is expressed as *τ*_*d*_ ≥ 0.5 and *τ*_*h*_ ≤ 0.5; i.e., transmission is higher to disease-affected than healthy individuals. We do not consider healthy individuals, as only disease-affected individuals are assumed to be sequenced, but we similarly restrict our subtype-specific transmission probabilities to be greater than or equal to 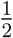. Since subtype *A* is assumed to more heritable than subtype *B* we impose the additional restriction that *τ*_*a*_ *> τ*_*b*_. Thus, under our alternative hypothesis the parameter space is 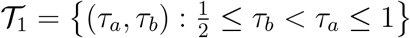, as illustrated in Figure 2.

**Figure 2:**
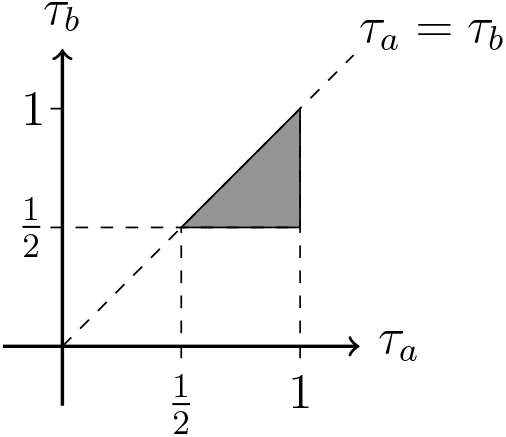
Alternative parameter space 𝒯 _1_.

### 2.2 Likelihoods

The above probability model leads to following likelihoods for the transmission parameters of interest, *τ*_*a*_ and *τ*_*b*_.

#### 2.2.1 Global

Each rare variant under consideration must have been observed in at least one disease-affected subject in the study; i.e., 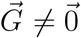. The likelihood is therefore 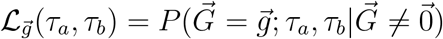.

Assuming independent families 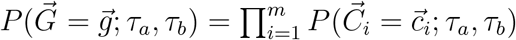, and, for 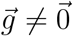,

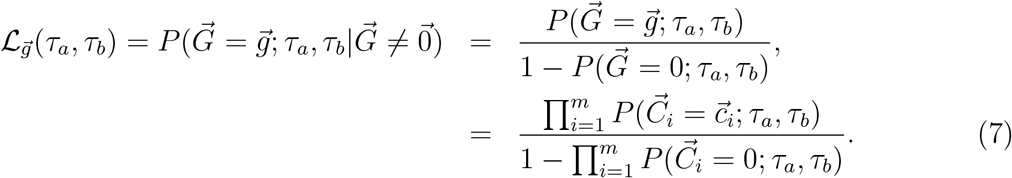

When a study contains only one pedigree, i.e. *m* = 1, and when 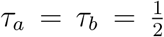 equation (7) is the RVS probability introduced by Bureau et al. [2014a]. However, when *m >* 1 the incorporation of individual familial configurations that may be identically 0 departs from Bureau et al. [2014a].

#### 2.2.2 Approximate unconditional

An approximate unconditional likelihood can be derived as follows. From equation (5), we can bound the probabilities of zero configurations, 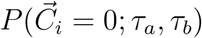, to be between 1 − *p*_*c*_*n*_*f*,*i*_ and one, where *p*_*c*_ is the population probability of carrying any of the cRVs and *n*_*f*,*i*_ is the number of founders in the the *i*th pedigree. When *p*_*c*_*n*_*f*,*i*_ is small, we can take 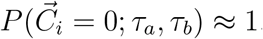. The *unconditional* global probability is then

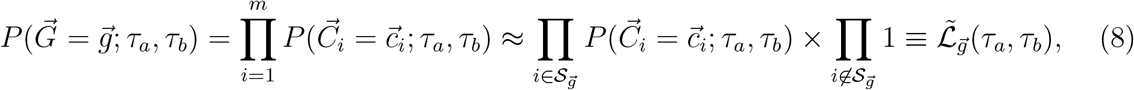

where 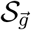 is the set of indices for the families with non-zero configurations. We call 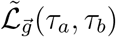 the approximate unconditional likelihood. To work well, *p*_*c*_*n*_*f*,*i*_ needs to be small (e.g. *n*_*f*,*i*_*p*_*c*_ *<* 0.01), such as with a very small carrier probability *p*_*c*_ and pedigrees with a modest number of founders.

#### 2.2.3 Local

An alternative to the above *global* approaches is a *local* approach in the spirit of Bureau et al. [2014b], that considers only the families with non-zero configurations (i.e. in 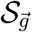). The local likelihood is

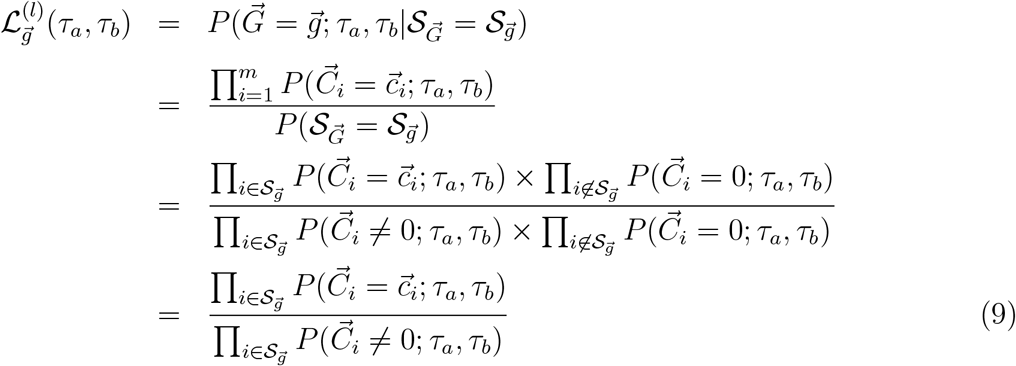

In the last line of equation (9), both the numerator, given by equation (4), and the denominator, given by equation (6), depend on *p*_*c*_ only through the common factor 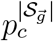, where 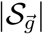 is the number of non-zero configurations. Hence the local likelihood does not depend on *p*_*c*_.

### 2.3 Statistics

The likelihoods defined above are used to form likelihood-ratio (LR) statistics for ranking RVs and testing the null hypothesis of no linkage and no association between an RV and disease subtypes. In this section we discuss these LR statistics and their p-values for testing. We also review the RVS statistic of Bureau et al. [2014b] and propose a modification that accounts for disease subtype.

#### 2.3.1 Likelihood ratios

Throughout, the maxima for LR statistics are obtained through a grid search over values of (*τ*_*a*_, *τ*_*b*_).

##### Global

The global LR statistic is based on the global likelihood 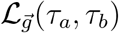 of equation (7). Letting 𝒯 = 𝒯 _0_ ∪ 𝒯 _1_ denote the parameter space, the global LR statistic is

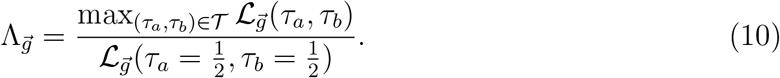

To understand how the global LR statistic depends on the carrier probability *p*_*c*_ we factor the global likelihood as

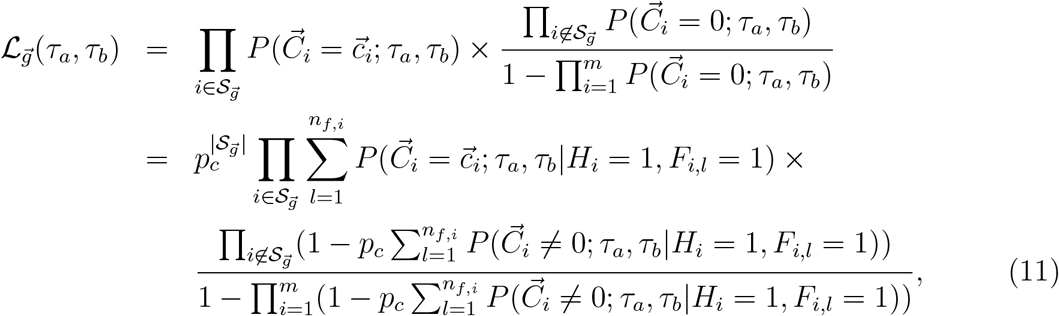

where the second line of the equation follows from inserting the expression for 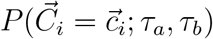 when 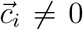 and 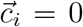 from equations (4) and (5), respectively, into the right-hand side of the first line. In the global likelihood ratio of equation (10), the factor 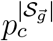 cancels out but *p*_*c*_ remains through the probabilities of familial configurations of zero. We return to this point in the Discussion.

For testing 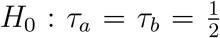, the distribution of global configurations 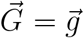 under the null hypothesis is the null distribution 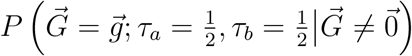. The p-value for an observed global configuration 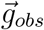 is then

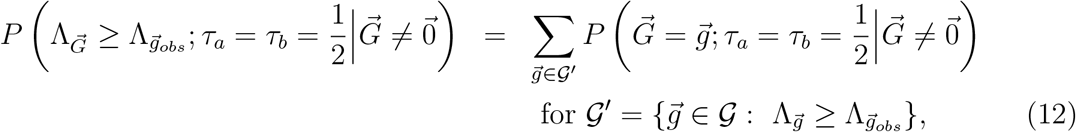

The sum on the right-hand side of the equation is over the set 𝒢 ′ of global configurations 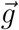 with global likelihood-ratio statistic as large as or larger than the observed value. This set is denoted 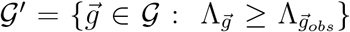, where 𝒢 is the sample space of non-zero global configurations.

##### Approximate unconditional

The approximate unconditional global likelihood-ratio statistic, or unconditional global LR statistic for short, is based on the approximate unconditional likelihood 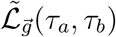 in equation (8):

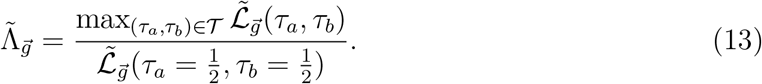

The null distribution for testing hypotheses is 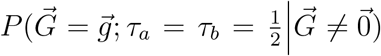, the same as for the global LR test. The p-value for an observed global configuration 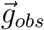 is thus the analog to equation (12) with 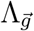 replaced by 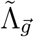. The unconditional global LR statistic does not depend on the carrier probability *p*_*c*_. Inserting the expression for 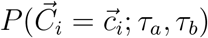 when 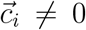 from equation (4) into equation (8) for the approximate unconditional likelihood yields an expression that depends on *p*_*c*_ only through the factor 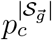. This factor cancels out of the likelihood ratio in equation (13). Thus, ranking with the unconditional global LR statistic does not depend on *p*_*c*_. By contrast, the null distribution *does* depend on *p*_*c*_, and hence so do p-values.

##### Local

The local LR statistic is formed from the local likelihood in equation (9):

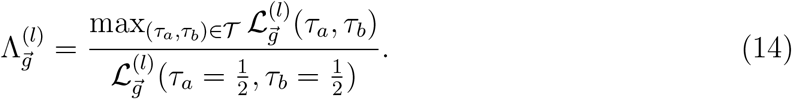

The null distribution for the local LR statistic is 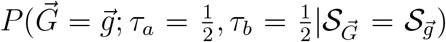. Since these local probabilities do not depend on *p*_*c*_ (see Section 2.2.3), neither does the local LR statistic nor its null probabilities. Therefore, ranking and testing based on the local LR statistic do not depend on *p*_*c*_.

#### 2.3.2 RVS methods

We considered ranking and testing using rare-variant sharing (RVS) approaches [Bureau et al., 2014a,b, 2019]. These methods use exact probabilities to test excess sharing of RVs observed in disease-affected relatives. The original RVS test was not developed for disease subtypes but is an established method that we include for comparison. We also propose an extension to account for disease subtypes.

##### Original

Without disease subtypes there is a single transmission parameter, *τ*. The null hypothesis for the test is 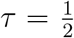 and the one-sided alternative hypothesis is that 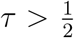. As defined in Bureau et al. [2019] the p-value of observed configuration 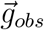 is given by:

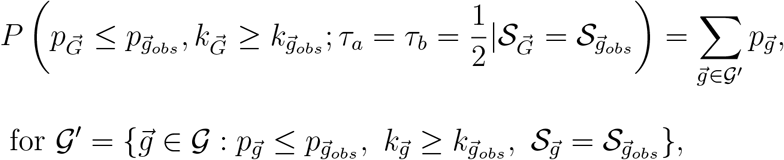

where 𝒢 is the sample space of 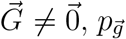 is RVS sharing probability for configuration 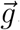, and 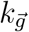 is the total number of sequenced subjects who share a variant in configuration 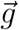. We use the p-value for testing and its inverse to rank configurations.

##### Modified

With disease subtypes there are two transmission parameters *τ*_*a*_ and *τ*_*b*_. The null hypothesis is 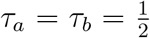 and, for the modified RVS test, the alternative hypothesis is that 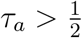 or 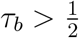. To accommodate disease subtype, we introduce an additional statistic *k*_*sub*_ for the number of subjects who carry the RV and are affected by the more heritable subtype. In the modified RVS test, the p-value of observed local configuration 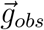 is:

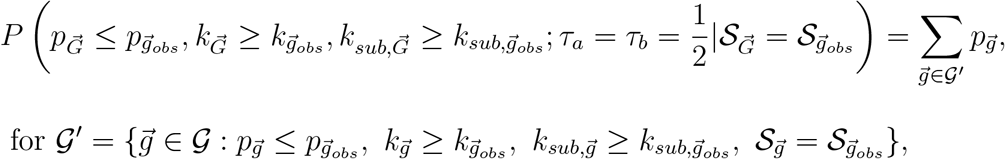

where 𝒢 is the sample space of 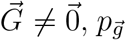 is RVS sharing probability for configuration 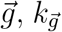 is the total number of sequenced subjects who share a variant in configuration 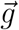, and 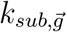 is the total number of sequenced subjects with the more heritable subtype who share a variant in configuration 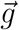. We use the p-value for testing and its inverse for ranking. The modified RVS approach incorporates the subtype of disease-affected relatives. However, unlike the transmission-based statistics presented earlier, this sharing approach does not consider the individual transmission events within families, which may provide additional information for prioritizing RVs.

To recap, this section on the statistics considered presents three LR methods and two RVS methods for ranking and testing rare variants. Of the LR methods, two are global and one is local, while the two RVS methods are both local. We note that the two global methods (global LR and unconditional global LR) condition on a less restrictive event that the RV is observed (or 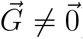) than the two local methods which condition on the event that the RV is observed in specific families (or 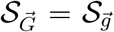). The less restrictive conditioning required for the global methods has the potential to improve power but requires specifying the carrier probability, *p*_*c*_, a nuisance parameter that is often unknown. However, as the specified carrier probability remains the same under the null and alternative hypotheses, the two global methods may be robust to misspecification of the nuisance parameter.

### 2.4 Simulation Methods

The goal of the simulation study was to assess ranking ability and power of the methods when *c*RVs are associated with disease status, and to verify that the methods maintain Type-I error rates at a nominal level when there are no *c*RVs. Ranking, power and Type-I error rate estimates are based on 2000 simulated studies. Briefly, a study is comprised of three simulated pedigrees ascertained according to criteria given below. For each pedigree, we sample founder sequences and use gene-dropping conditional on *c*RV status from the founders to the sequenced affected individuals. We then use each of the five methods (global LR, unconditional global LR, local LR, RVS and modified RVS) to (i) rank the observed *c*RVs against the other RVs observed in the sequenced affected individuals and (ii) test the observed *c*RVs for association with disease status. Additional details on each of the major steps are given in the following subsections.

#### 2.4.1 Simulated pedigrees

Motivated by the Lymphoid Cancer Families Study, we simulated pedigrees segregating *c*RVs using the `SimRVPedigree` [Nieuwoudt et al., 2018] R package [R Core Team, 2017]. Pedigrees contained two disease subtypes, labelled Hodgkin Lymphoma (HL) and non-Hodgkin Lymphoma (NHL), with HL considered to be the more heritable subtype. Genetic relative risks (GRRs) for HL and NHL in *c*RV carriers were set to 35 and 1, respectively. The *c*RV carrier probability was set to 0.00032. Disease onset and death for individuals in the pedigree were simulated with the age-specific hazard rates for lymphoid cancer provided in the `SimRVPedigree` package. In the package, baseline and disease-subtype-specific hazard rates for onset of and death from HL and NHL are based on the Surveillance, Epidemiology, and End Results (SEER) database [Surveillance, Epidemiology, and End Results (SEER) Program, a,b,c]. Hazards rate for death in unaffected individuals are based on general population data [Bell and Miller, 2005]. Pedigrees were retained if they included four to six known disease-affected relatives, with at least one of each disease subtype. Pedigrees with more than six disease-affected relatives were rare and excluded to keep computations manageable. For simulations under the alternative hypothesis pedigrees were also required to include at least one HL case that carried a *c*RV. Ascertaining for at least four affected members is a strategy to increase the chance of a genetic cause for the disease in a pedigree, although in practice such pedigrees may be difficult to find. Further details on the simulation settings are given in Appendix C.6 of Nieuwoudt [2021].

As *de novo* simulation of ascertained pedigrees is time consuming, we created a pool of 55 ascertained pedigrees in advance and then sampled three of these, without replacement, from the pool to make a study. A study size of three pedigrees is reasonable when families with four or more affected members are uncommon in the population as is the case for lymphoid cancer [Jones et al., 2017]. The small study size also keeps computation times manageable for the analysis of the simulated datasets. Section 4 provides further discussion of the computational limitations of the methods. Appendix A shows that it is feasible to rank all variants in a much larger study of all 55 ascertained families in the pool.

#### 2.4.2 Population of founder sequences

Founder sequences are drawn from a population of simulated exome sequences containing single-nucleotide variants with population minor allele frequency (MAF) < 0.01, available at *https://zenodo.org/record/6369360* [Epasinghege Dona and Graham, 2022]. Briefly, exome sequences were generated by the evolutionary simulation program SLiM [Haller and Messer, 2017], under the “American Admixed” demographic model of the `stdpopsim` library [Adrion et al., 2020], a mutation model in which the first two base pairs of a codon are under negative selection and the third base pair is selectively neutral, and a recombination model following the recombination map in the `SimRVSequences` R package [Nieuwoudt et al., 2019]. From this data source we selected chromosome 8 and further filtered to variants with MAF < 0.001.

We use the `SimRVSequences` package to “gene drop” founder sequences through simulated pedigrees conditional on *c*RV status. The package requires the specification of a set of candidate *c*RVs. We chose ten *c*RVs from the genes *TNFRSF10B* and *TNFRSF11B* on chromosome 8 in the human apoptosis pathway as follows. From these genes, ten RVs with population minor allele count ≤ 10 were sampled in proportion to their selection coefficients, yielding four singletons, five doubletons and one tripleton with a cumulative population frequency of 0.0001578. The implied carrier probability of a *c*RV is therefore about 2 × 0.0001578 = 0.00032. Table 1 summarizes the *c*RVs.

**Table 1:** Chromosome, gene, base-pair position, minor allele frequency (MAF) and selection coefficient of the 10 RVs randomly chosen to be causal in the simulations under the alternative hypothesis.

| chrom | gene | position | MAF | selection coefficient |
| --- | --- | --- | --- | --- |
| 8 | *TNFRSR10B* | 23020319 | 0.000009 | -0.009173 |
| 8 | *TNFRSR10B* | 23020346 | 0.000009 | -0.000410 |
| 8 | *TNFRSR10B* | 23020454 | 0.000019 | -0.006706 |
| 8 | *TNFRSR10B* | 23021159 | 0.000028 | -0.023490 |
| 8 | *TNFRSR10B* | 23021957 | 0.000009 | -0.001670 |
| 8 | *TNFRSR10B* | 23022671 | 0.000009 | -0.000658 |
| 8 | *TNFRSR10B* | 23068278 | 0.000019 | -0.012389 |
| 8 | *TNFRSR10B* | 23068413 | 0.000019 | -0.001102 |
| 8 | *TNFRSF11B* | 118923860 | 0.000019 | -0.008192 |
| 8 | *TNFRSF11B* | 118933213 | 0.000019 | -0.017692 |

The specifics of gene-dropping sequences through pedigrees and the subsequent ranking and testing of *c*RVs depend on whether simulations are under the alternative or the null hypothesis, as described next.

#### 2.4.3 Alternative hypothesis

##### Conditional gene-drop

Data for the sequenced affected individuals in a study is obtained with `SimRVSequences`. For each pedigree, a *c*RVs is sampled from the list of candidates and then founder sequences are sampled from the population and dropped through the pedigree conditional on *c*RV status. The resulting sequence data from the study is then filtered to variants that are observed in at least one sequenced affected individual. In terms of the notation of Section 2.1 these are the variants with global configuration 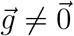. As *c*RVs are sampled with replacement from a list of candidates, the same *c*RV may be sampled for multiple pedigrees. Hence a study can have one, two or three *c*RVs.

##### Ranking

We determine the rank of a given *c*RV relative to all RVs observed in the study. In our ranking system, suppose *N*_*tot*_ RVs have been observed among the *m* pedigrees in the study and that the *i*^*th*^ RV has statistic value *w*_*i*_ for *i* ∈ {1, 2, …, *N*_*tot*_}. Since studies yield slightly different numbers of RVs, we normalize the ranks to compare them across studies.

The normalized rank of the *i*^*th*^ statistic is

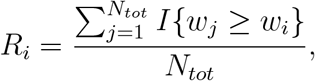

where *I*{*w*_*j*_ ≥ *w*_*i*_} = 1 if *w*_*j*_ ≥ *w*_*i*_, and 0 otherwise. Normalized ranks are ∈ (0, 1], with values closer to zero indicating top ranks and those closer to one indicating bottom ranks. The unnormalized, raw ranks are the sample ranks in reverse order, with ranks for tied values assigned the maximum rank [R Core Team, 2017]. For a given method, we summarize the distribution of *c*RV ranks over the 2000 simulated studies by the median and inter-quartile range (IQR). Recall that there can be one, two or three *c*RVs for a given study. We compare the distributions of (i) the average *c*RV rank for each study, (ii) the top-ranked *c*RV for each study, (iii) the second-ranked *c*RV for each study and (iv) the third-ranked *c*RV for each study. If a study has only one *c*RV, its average is based on a single observation.

##### Power

The estimated power for a given method is the proportion of all *c*RV tests, over all simulated studies and all *c*RVs, that reject the null hypothesis at the 5% level.

#### 2.4.4 Null hypothesis

As no *c*RVs are present under the null hypothesis, we do not rank RVs. We do however test RVs to estimate Type-I error rates.

##### Unconditional gene-drop

Under the null hypothesis, we simulate sequence data by random sampling of founder sequences and *unconditional* gene-dropping through the pedigrees. In principle, every RV observed in this way is “null” and could be tested for association with disease status to estimate the Type-I error rates. However, to match the target variant frequencies with those under the alternative hypotheses we chose to repeat the gene-drop simulation until at least one of the candidate *c*RVs (under the alternative hypothesis) was observed in the study.

##### Type-I error

All candidate *c*RVs observed in the unconditional gene-drop simulations are tested at the 5% level. The estimated Type-I error rate for a given method is the proportion of all such tests, over all simulated studies and all candidate *c*RVs, that reject the null hypothesis at the 5% level.

## 3 Results

### 3.1 Example data analysis

To illustrate the data and methods, we present an example analysis of a hypothetical study containing three families simulated under the alternative hypothesis, using the methods outlined in Section 2.4. The families for this hypothetical study are shown in Figures 1, 3 and 4. All families have been “trimmed” to include only the affected members and their ancestors back to the founders.

**Figure 3:**
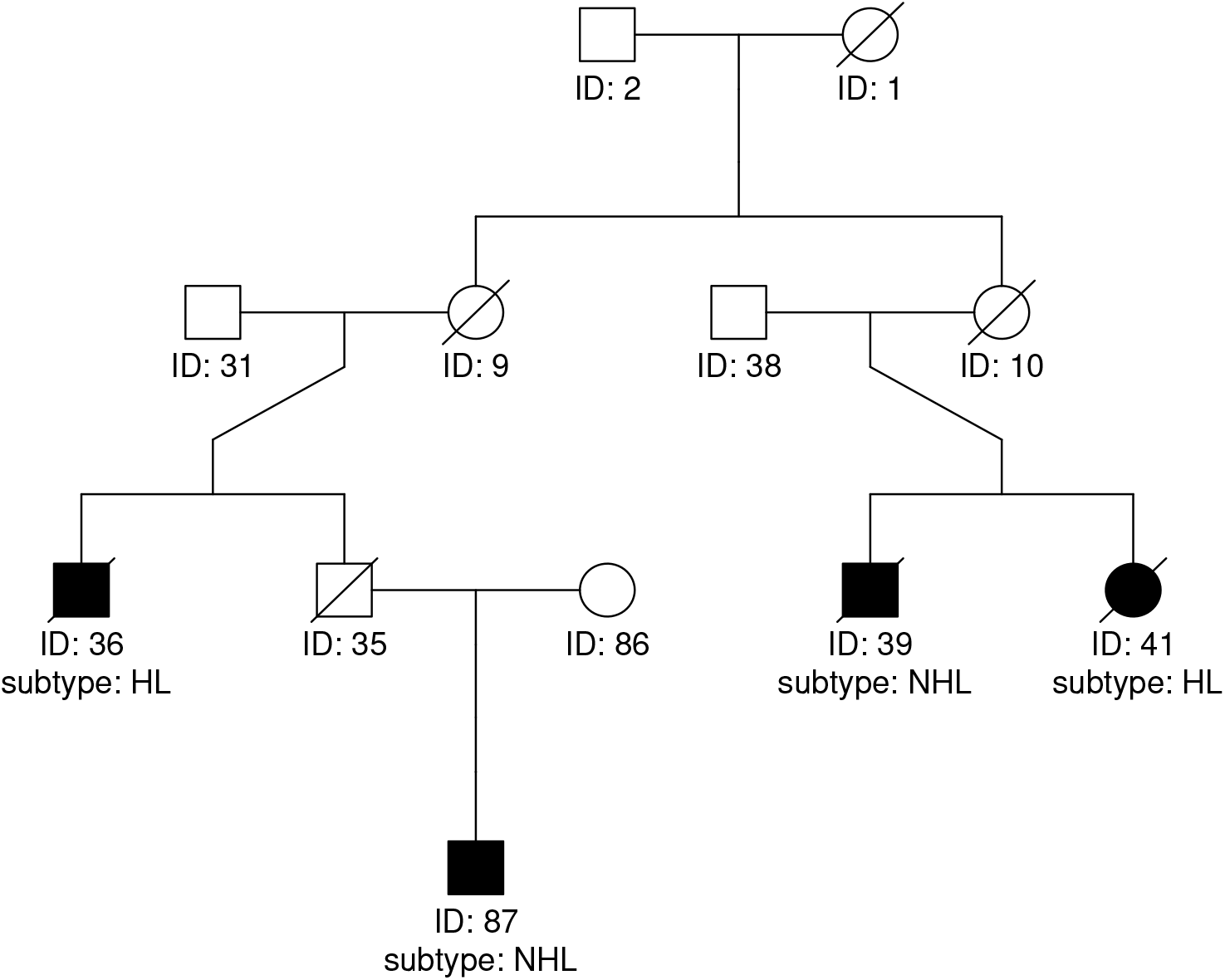
Second simulated example family. Affected members have black symbols, with disease subtype given below. Members with known date of death have a slash through their symbol.

**Figure 4:**
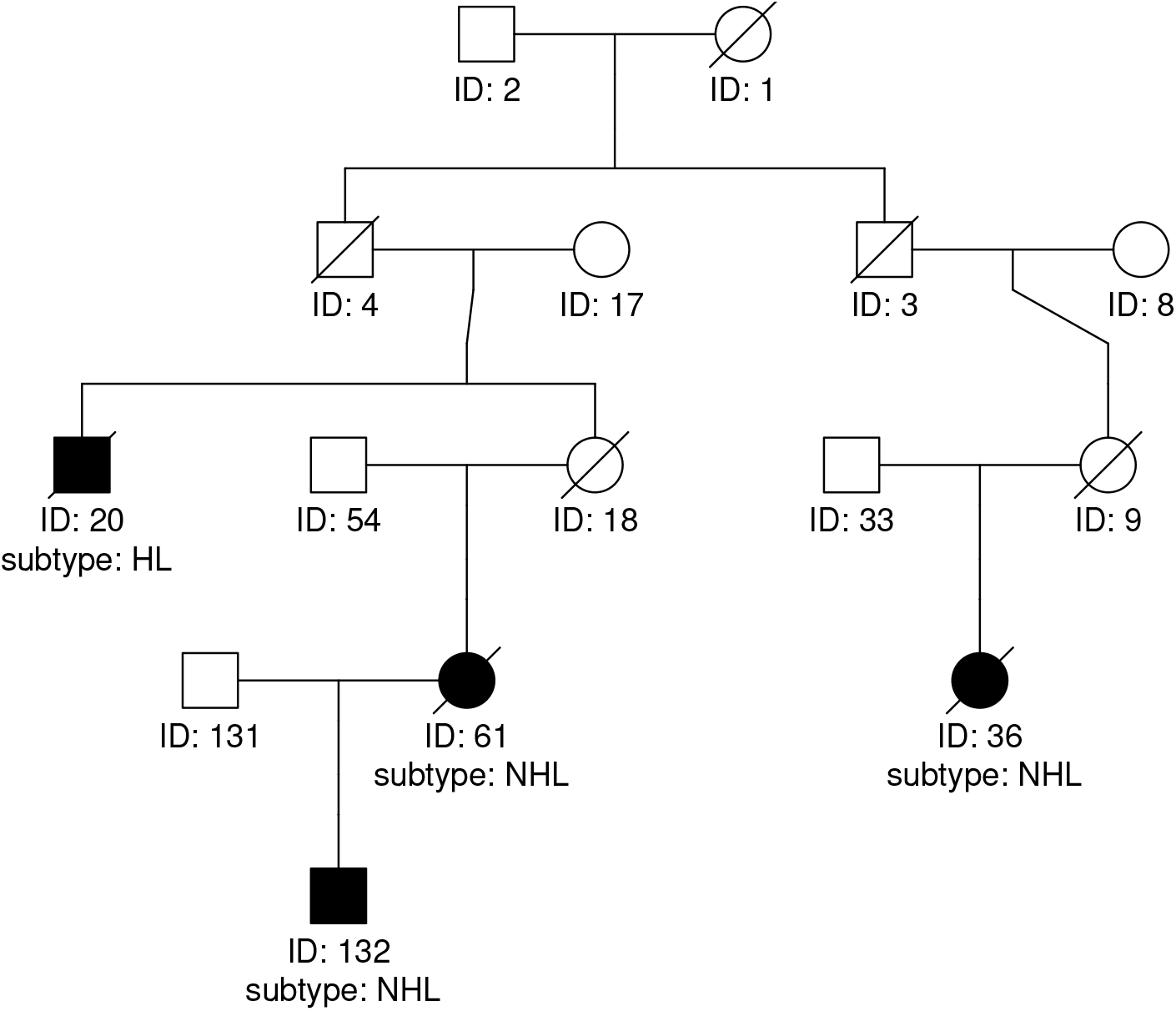
Third simulated example family. Affected members have black symbols, with disease subtype given below. Members with known date of death have a slash through their symbol.

#### 3.1.1 Study summaries

The chromosome-8 exome sequences of the affected individuals contain 74 RVs. Two of these RVs appear in more than one founder of a given pedigree, and so are inconsistent with the assumption that a *c*RV enters a pedigree through at most one founder. After removing these two RVs, 72 RVs remain for the analysis. Of the remaining 72 RVs, 57 appear only once, 13 appear twice and two appear three times. Table 2 summarizes the counts of RVs that appear more than once, by pedigree and disease subtype.

**Table 2:** Counts of RVs within each pedigree by disease subtype, for RVs that appear in more than one affected individual in the study. Only the RV 8_6708389 appears in more than one pedigree.

|  |  |  | ped 1 |  | ped 2 |  | ped 3 |  |
| --- | --- | --- | --- | --- | --- | --- | --- | --- |
|  |  |  | NHL | HL | NHL | HL | NHL | HL |
| Number of affected individuals: |  |  | 2 | 2 | 2 | 2 | 3 | 1 |
| RV | chrom | position |  |  |  |  |  |  |
| 8_6708389 | 8 | 6708389 | 1 | 0 | 0 | 1 | 0 | 0 |
| 8_7972232 | 8 | 7972232 | 0 | 0 | 0 | 2 | 0 | 0 |
| 8_8376739 | 8 | 8376739 | 1 | 1 | 0 | 0 | 0 | 0 |
| 8_12116292 | 8 | 12116292 | 0 | 0 | 2 | 0 | 0 | 0 |
| 8_23020454 | 8 | 23020454 | 0 | 2 | 0 | 0 | 0 | 0 |
| 8_23068413 | 8 | 23068413 | 0 | 0 | 0 | 2 | 0 | 0 |
| 8_30846116 | 8 | 30846116 | 0 | 0 | 2 | 0 | 0 | 0 |
| 8_39680176 | 8 | 39680176 | 1 | 2 | 0 | 0 | 0 | 0 |
| 8_41261959 | 8 | 41261959 | 0 | 0 | 2 | 0 | 0 | 0 |
| 8_54047456 | 8 | 54047456 | 0 | 0 | 1 | 1 | 0 | 0 |
| 8_60866956 | 8 | 60866956 | 1 | 1 | 0 | 0 | 0 | 0 |
| 8_65161224 | 8 | 65161224 | 0 | 0 | 0 | 0 | 2 | 1 |
| 8_103382957 | 8 | 103382957 | 1 | 1 | 0 | 0 | 0 | 0 |
| 8_107249922 | 8 | 107249922 | 0 | 0 | 1 | 1 | 0 | 0 |
| 8_142271244 | 8 | 142271244 | 0 | 0 | 1 | 1 | 0 | 0 |

#### 3.1.2 Ranking

We calculate the raw ranks of the 72 RVs being analysed. The top-10 RVs by each ranking method are shown in Table 3. The ranks of the three *c*RVs in the study are shown in Table 4. We see that the *c*RV 8_23020454 from pedigree 1 is ranked second by the LR-based methods and third by the RVS-based methods. This *c*RV appears in the two HL-affected individuals in the pedigree, who are second cousins. The *c*RV 8_23068413 from pedigree 2 is ranked fourth by the LR-based methods and ninth by the RVS-based methods. This *c*RV appears in the two HL-affected individuals in the pedigree, who are first cousins and therefore separated by fewer meioses than the second cousins in pedigree 1. The *c*RV 8_118923860 from pedigree 3 has rank anywhere from 15 to 49 out of 72. This *c*RV was observed in only one HL-affected individual.

**Table 3:** Top-10-ranked RVs for the different ranking methods. The ranks from the global LR methods with assumed carrier probabilities of 1/100, 1/10, 1/2, 1, 2 and 10 times the true value of 0.00032 are all identical and given in the column labelled gLR1to6. Rankings from the global LR method with assumed carrier probability of 100 times the true value are in the column labelled gLR7. The other ranking methods such as the unconditional global LR (ungLR), local LR (loLR), original RVS (RVS) and modified RVS (mRVS), do not require a value of the carrier probability. The *c*RVs 8_23020454 and 8_23068413 from the first and second pedigree, respectively, are in bold. The *c*RV 8_118923860 from the third pedigree was not ranked in the top 10 by any method.

| raw<br>rank | Method |  |  |  |  |  |
| --- | --- | --- | --- | --- | --- | --- |
|  | gLR1to6 | gLR7 | ungLR | loLR | RVS | mRVS |
| 1 | 8_39680176 | 8_39680176 | 8_39680176 | 8_39680176 | 8_65161224 | 8_39680176 |
| 2 | **8_23020454** | **8_23020454** | **8_23020454** | **8_23020454** | 8_39680176 | 8_65161224 |
| 3 | 8_7972232 | 8_7972232 | 8_7972232 | 8_7972232 | **8_23020454** | **8_23020454** |
| 4 | **8_23068413** | **8_23068413** | **8_23068413** | **8_23068413** | 8_8376739 | 8_8376739 |
| 5 | 8_65161224 | 8_65161224 | 8_65161224 | 8_65161224 | 8_12116292 | 8_12116292 |
| 6 | 8_8376739 | 8_8376739 | 8_8376739 | 8_8376739 | 8_30846116 | 8_30846116 |
| 7 | 8_54047456 | 8_60866956 | 8_54047456 | 8_54047456 | 8_41261959 | 8_41261959 |
| 8 | 8_107249922 | 8_103382957 | 8_107249922 | 8_107249922 | 8_7972232 | 8_7972232 |
| 9 | 8_60866956 | 8_54047456 | 8_60866956 | 8_60866956 | **8_23068413** | **8_23068413** |
| 10 | 8_103382957 | 8_107249922 | 8_103382957 | 8_103382957 | 8_60866956 | 8_60866956 |

**Table 4:**
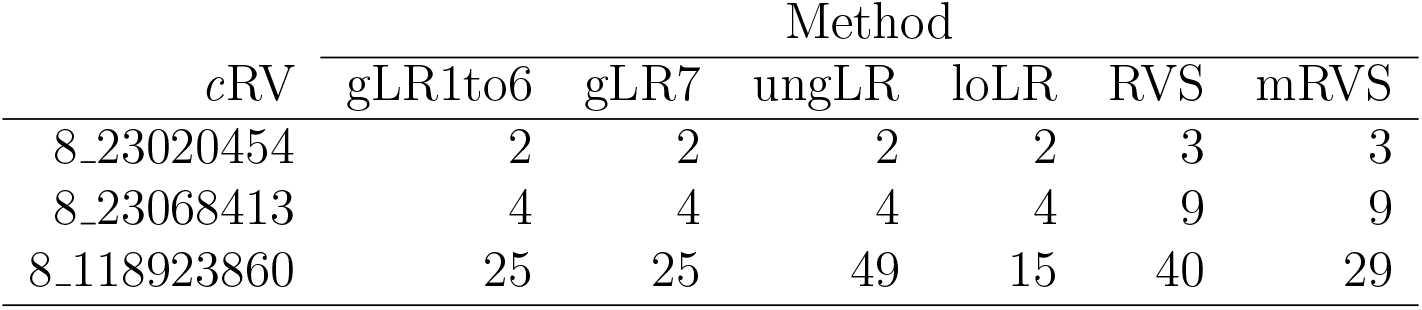
Raw ranks of the three *c*RVs in the study. The ranks from the global LR methods with carrier probabilities assumed to be 1/100, 1/10, 1/2, 1, 2 and 10 times the true value of 0.0032 are all identical and given in the column labelled *gLR1to6*. Rankings from the global LR method with carrier probability assumed to be 100 times the true value are in the column labelled `gLR7`. The other ranking methods do not depend on a value of the carrier probability.

We also note the similarity of rankings produced by the three LR methods. For example, as shown in Table 3, the global LR, unconditional global LR and local LR methods have the same top-10 rankings once gLR7, the global LR method with largest assumed carrier probability is excluded. The gLR7 method assumes an unrealistically large carrier probability of 0.032, or 100 times the true value. Similar results for the top-10 rankings hold in the other simulated datasets as well (results not shown).

#### 3.1.3 Testing

Table 5 shows p-values from tests of association between the three study *c*RVs and disease status for the different testing methods. The *c*RVs 8_23020454 from pedigree 1 and 8_23068413 from pedigree 2 were observed in two HL-affected individuals and have reasonably small p-values (e.g., 0.0056 and 0.0174, respectively, by the global methods). The cRV 8_118923860 from pedigree 3 was observed in only one HL-affected individual and has large p-values (*>* 0.31).

**Table 5:**
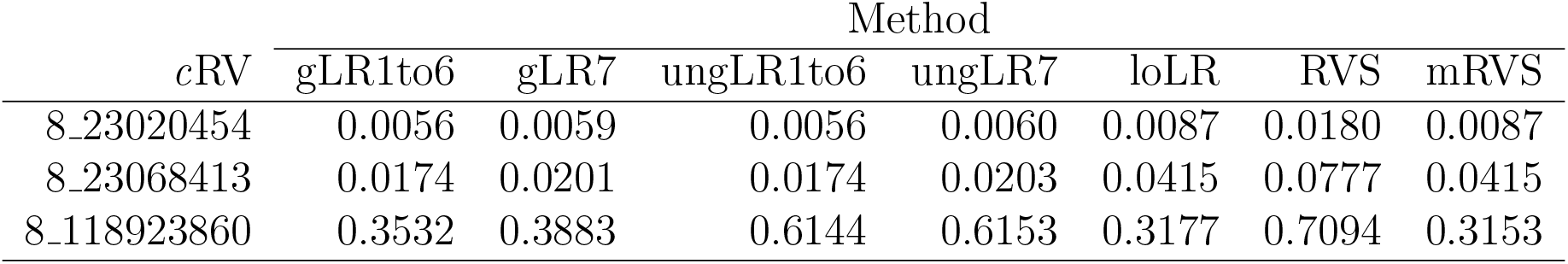
Results of association tests for the three study *c*RVs from the different methods. The p-values from the global LR and unconditional global LR methods with assumed carrier probabilities of 1/100, 1/10, 1/2, 1, 2 and 10 times the true value of 0.0032 are the same to four decimals and reported in the columns labelled `gLR1to6` and `ungLR1to6`, respectively. The p-values from the global LR and unconditional global LR tests with assumed carrier probability of 100 times the true value are reported in the columns labelled `gLR7` and `ungLR7`, respectively. The other testing methods do not require a value of the carrier probability.

### 3.2 Simulation study

Of the 2000 studies simulated under the alternative hypothesis, 34 contained a single RV from the list of ten candidate *c*RVs, 632 contained two RVs from the candidate list, and 1334 contained three RVs from the list. Of the 2000 studies simulated under the null hypothesis, 1996 contained a single RV from the list of candidate *c*RVs and four contained two RVs from the candidate list.

#### 3.2.1 Ranking

The total number of RVs per study varied from 48 to 132, with median value 84. To compare the *c*RV ranks across studies, we normalized them by dividing by the total number of RVs in the study. For example, in a study with 84 RVs, a *c*RV with raw rank of 10 has a normalized rank of 10/84 ≈ 0.12 whereas in a study with 48 RVs, it has a normalized rank of 10/48 ≈ .21. Figure 5 shows a graphical summary of the normalized ranking results for the different methods. As rankings for the global LR method with *p*_*c*_ set to 1/100, 1/10, 1/2, 1, 2 and 10 times the true value were very similar (results not shown), they are represented on the plot by the global LR ranking with true value, *p*_*c*_ = 0.00032.

**Figure 5:**
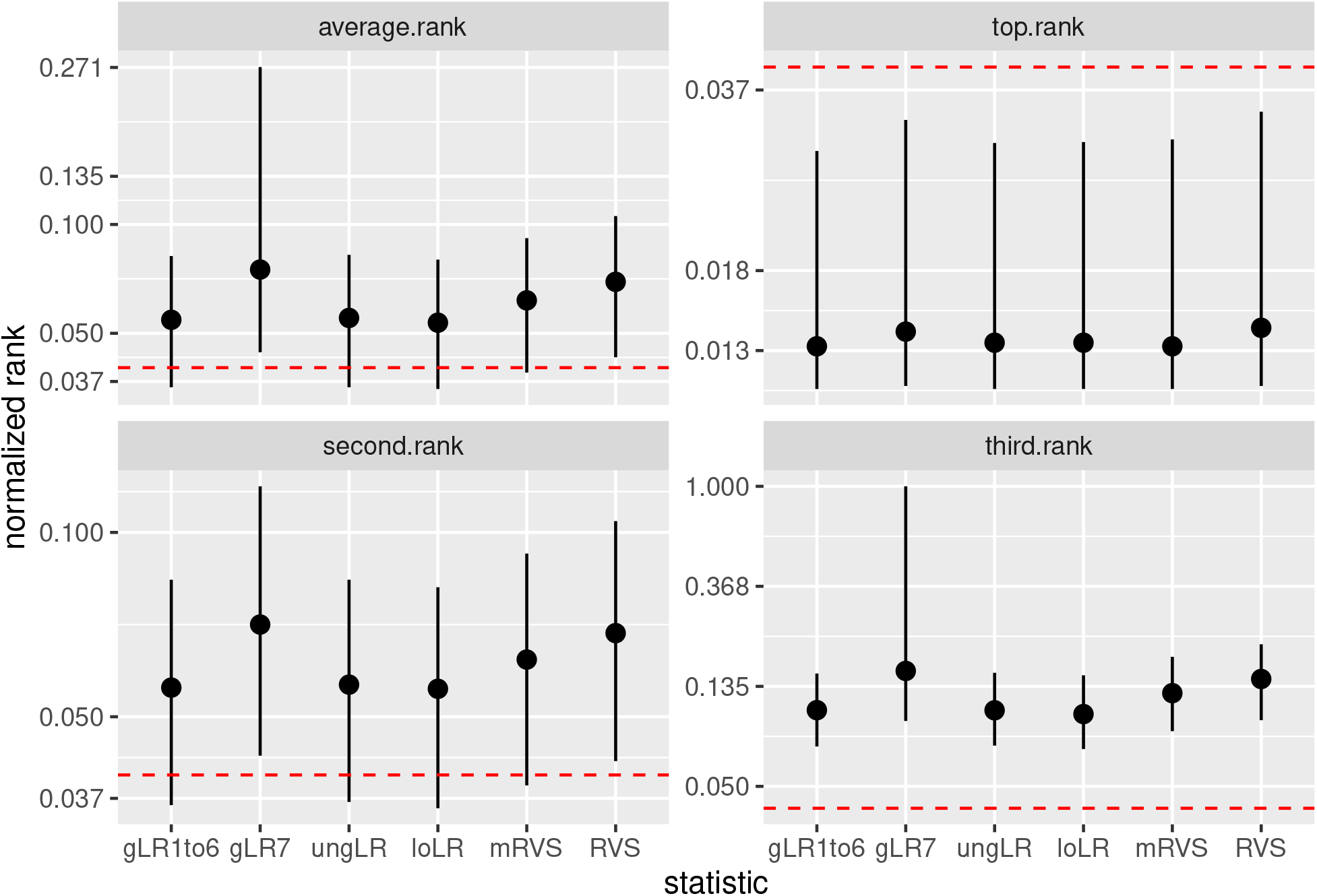
Median normalized ranks (dots) and IQR (bars) from the different methods using the average (top left), first (top right), second (bottom left) or third (bottom right) rank of the *c*RVs in each study. The methods are labelled as: GLR1to6, global LR statistic with values of *p*_*c*_ set to 1/100, 1/10, 1/2, 1, 2 or 10 times the true value; gLR7, global LR statistic with *p*_*c*_ set to 100 times the true value; ungLR, unconditional global LR; loLR, local LR; mRVS, modified RVS; or RVS. Summary statistics are plotted on the log scale, but the y-axis labels are on the original scale of the normalized ranks. The red horizontal dashed line indicates a normalized rank of 0.04.

The *c*RV rankings from the global LR statistic are similar across the specified values of the carrier probability *p*_*c*_, except for when *p*_*c*_ is misspecified to be 100 times its true value (results not shown). The distributions of the *c*RV rankings from the global LR and unconditional global LR statistics are also similar, except when the value of the carrier probability is misspecified to be 100 times its true value. The local LR statistic tends to rank the *c*RVs as high as or higher than the global approaches as indicated by its slightly lower median rank. The LR-based statistics tend to rank the *c*RVs higher than the RVS and modified RVS statistics, as indicated by their smaller median ranks, except for the global LR method with carrier probability misspecified to be 100 times its true value. As expected, the RVS approach tends to rank *c*RVs the lowest among the methods because it was not developed with disease subtypes in mind. The modified RVS approach accounts for disease subtypes and tends to rank *c*RVs higher than the naive RVS approach but not as highly as the likelihood approaches.

#### 3.2.2 Type I error rates

The estimated Type-I error rates for the different methods are shown in Table 6. Estimates are based on 2004 RVs from the list of candidate *c*RVs sampled under the null hypothesis. All tests control the Type-I error rate at the nominal 0.05 level, and in fact all but the modified RVS appear to be conservative. In particular, the LR tests (local LR, global LR and unconditional global LR) and the RVS test have estimated Type-I error rates at 3 or more SEs below 0.05.

**Table 6:** Estimated Type-I error rate (SE), all tested RVs (2004 in total)

| Factor* | Method |  |  |  |  |
| --- | --- | --- | --- | --- | --- |
|  | Local |  |  | Global |  |
|  | RVS | mRVS | loLR | gLR | ungLR |
| NA | 0.041 (0.003) | 0.052 (0.003) | 0.028 (0.002) | — | — |
| 1/100 | — | — | — | 0.037 (0.002) | 0.035 (0.002) |
| 1/10 | — | — | — | 0.037 (0.002) | 0.035 (0.002) |
| 1/2 | — | — | — | 0.036 (0.002) | 0.035 (0.002) |
| 1 | — | — | — | 0.036 (0.002) | 0.034 (0.002) |
| 2 | — | — | — | 0.036 (0.002) | 0.034 (0.002) |
| 10 | — | — | — | 0.033 (0.002) | 0.033 (0.002) |
| 100 | — | — | — | 0.007 (0.001) | 0.024 (0.002) |

\* Assumed carrier probability is factor times true value $p_c = 0.000320$ .

#### 3.2.3 Power

Recall that in our 2000 simulated studies 1334 studies had three *c*RVs, 632 studies had two *c*RVs and 34 studies had one sampled *c*RV, for a total of 5300 *c*RVs considered across all studies. The resulting estimates of power are shown in Table 7.

**Table 7:** Estimated power (SE), all cRVs (5300 assessed in total)

| Factor* | Method |  |  |  |  |
| --- | --- | --- | --- | --- | --- |
|  | Local |  |  | Global |  |
|  | RVS | mRVS | loLR | gLR | ungLR |
| NA | 0.692 (0.006) | 0.759 (0.006) | 0.758 (0.006) | — | — |
| 1/100 | — | — | — | 0.835 (0.005) | 0.831 (0.005) |
| 1/10 | — | — | — | 0.834 (0.005) | 0.831 (0.005) |
| 1/2 | — | — | — | 0.832 (0.005) | 0.831 (0.005) |
| 1 | — | — | — | 0.832 (0.005) | 0.831 (0.005) |
| 2 | — | — | — | 0.832 (0.005) | 0.830 (0.005) |
| 10 | — | — | — | 0.822 (0.005) | 0.824 (0.005) |
| 100 | — | — | — | 0.429 (0.006) | 0.733 (0.006) |

\* Assumed carrier probability is factor times true value $p_c = 0.000320$ .

From Table 7, global methods appear to be robust to misspecification of the carrier probability, except when the carrier probability is substantially overspecified (i.e. more than 10 times larger than the true value). In particular, overspecifying the carrier probability to be 100 times too large leads to a conservative test with lower power, especially for the global LR test. The estimated power of the global LR and unconditional global LR tests are similar, and larger than the estimated power of the local tests (local LR, RVS and modified RVS), except when the value of the carrier probability is substantially overspecified. In particular, when the carrier probability is specified to be 100 times larger than its true value, the global LR test has lower power than the unconditional global LR test and both these global tests have lower power than the local tests.

## 4 Discussion

We have developed likelihood-based methods for prioritizing and testing rare variants shared by affected relatives in families containing multiple disease cases where one subtype is more heritable than the others. Global likelihood approaches condition on a variant being observed in the study and assume a known value of the population probability, *p*_*c*_, of carrying a causal variant. By contrast, the local likelihood approach conditions on a variant being observed in specific families to eliminate the unknown nuisance parameter *p*_*c*_ from the likelihood. We have also extended the RVS method of Bureau and colleagues [Bureau et al., 2014a,b, 2019] to accommodate disease subtypes. Through simulation we compared the different methods in terms of their ranking ability and power to detect *c*RVs.

In the presence of disease subtypes, our main conclusions are as follows. First, when one subtype is more heritable, all methods that consider disease subtype are able to rank and detect *c*RVs better than the original RVS method which is naïve to disease subtypes. When disease subtypes are not meaningful (i.e., subtypes are equally heritable), the proposed methods perform similarly to the original RVS method (results not shown). For studies that ascertain only one subtype of disease, the proposed methods with two transmission parameters *τ*_*a*_ and *τ*_*b*_ can be simplified to a *single* transmission parameter, *τ*, but such modifications would not be expected to offer an improvement over the RVS method.

For prioritizing RVs, the LR methods (global LR, unconditional global LR and local LR) generally rank better than the RVS methods, including the modified RVS approach that accounts for disease subtype. To understand this difference in ranking ability, consider the local LR and modified RVS, both of which are local methods that consider disease subtypes.

The statistics for the local LR and modified RVS methods use a restricted and less restricted form of the alternative hypothesis, respectively. Specifically, the local LR method incorporates prior information about the more heritable subtype through its restricted alternative hypothesis, *τ*_*a*_ *> τ*_*b*_ ≥ 1/2, while the modified RVS method uses a less restricted alternative hypothesis that *τ*_*a*_ or *τ*_*b*_ is *>* 1*/*2. Subtype labels matter under the alternative hypothesis of the local-LR but not the modified-RVS method. The consequences of this difference in alternative hypotheses can be illustrated with the *c*RVs 8_23020454 and 8_23068413 that were present in more than one sequenced affected individual in the example dataset (see Section 3.1). Swapping the disease subtype labels of sequenced affected individuals changes the local-LR rankings but not the modified-RVS rankings. In particular, the local-LR rankings of 8_23020454 and 8_23068413 change from 2 to 11 and from 4 to 44 respectively when the subtype labels are swapped, while the modified-RVS rankings are 3 and 9, respectively, regardless of subtype labels.

The generally better ranking of the LR methods has an exception. In our simulations, the ranking performance of the global LR statistic suffers when the carrier probability is over-specified by a factor of more than 10 times its true value. From equation (11), we see that the global likelihood ratio depends on the carrier probability, *p*_*c*_, only through the probabilities of familial configurations of zero. Familial configurations of zero can arise if: (i) a *c*RV was not introduced into the pedigree (no *c*RV), or (ii) a *c*RV was introduced into the pedigree through one of the founders but was not transmitted from the founder to any of the sequenced affected individuals (*c*RV not transmitted). A small value of *p*_*c*_ makes the no-*c*RV scenario more likely and the *c*RV-not-transmitted scenario less likely. Conversely, a large value of *p*_*c*_ makes the no-*c*RV scenario less likely and the *c*RV-not-transmitted scenario more likely, thereby attenuating estimates of the transmission parameters. Attenuated estimates of the transmission parameters give *c*RVs unremarkable global LR statistics, which makes *c*RV ranks more similar to the ranks of other RVs (and reduces power to detect *c*RVs). In contrast to the global LR statistic, the unconditional global LR and local LR statistics don’t depend on the carrier probability which makes these statistics more suitable for ranking and prioritizing RVs.

We also investigated the statistical properties of the tests for *c*RVs. As expected, all tests control the Type-I error rate at the nominal 5% level and, in fact, all but the modified RVS test were conservative. With respect to power, the global tests (global LR and unconditional global LR) have more power to detect *c*RVs than the local tests (local LR, modified RVS and RVS) for reasonable values of the assumed *c*RV carrier probability *p*_*c*_ (i.e. 10 times or lower than the true value) because local conditioning leads to a loss of information. The power of the global methods is similar. We also found that the global methods are robust to mis-specification of the carrier probability except when it is substantially overspecified. In particular, specifying the carrier probability to be 100 times too large leads to an overly conservative test with lower power, especially for the global LR test. As already noted in our discussion of the ranking results, the global LR test loses power because substantially over-specifying the carrier probability attenuates the estimates of the transmission probability parameters. In general, when using the global LR method, we recommend erring on the side of under-specifying the assumed carrier probability for both ranking and testing.

The RVS test was not developed to handle disease subtypes but is included here as a reference approach. To account for disease subtypes, we have extended the original RVS test. Both the original and modified RVS methods are “local”; i.e., include only families observed to carry the RV being tested. Among the local methods developed to handle more than one disease subtype, the local LR test is conservative whereas the modified RVS is not. We observed the local LR test to have similar power to the modified RVS test.

In principle, the likelihood-based approaches can be extended to more than two disease subtypes by adding additional subtype-specific transmission parameters to the model. However, even with just two parameters, these approaches are hampered by the limited number of transmissions within families and the incomplete information about transmission parameters encoded in familial configurations. This lack of information in the data can lead to unstable or even undefined parameter estimates [Nieuwoudt, 2021, Appendix C.3] and necessitates a simple model for transmission probabilities. When there are more than two subtypes, we recommend categorizing them into two groups based on heritability instead of introducing extra transmission parameters into the model.

Computation times for statistical testing of RVs grow exponentially while those for likelihood-based ranking grow linearly in the number of affected individuals (see Appendix A). As a result, testing is only feasible for relatively small studies involving a handful of families. By contrast, likelihood-based ranking allows researchers to prioritize RVs from much larger studies. For example, Appendix A presents a likelihood-based ranking analysis of all 55 families used in the simulation study. As RVS-based methods rely on local p-values to prioritize RVs, RVS-based ranking is not feasible for larger family studies.

The family-based association statistics we investigate aim to identify rare causal variants with large effects, which is also the motivation for traditional linkage analysis. Linkage analysis relies on identifying shared identical-by-descent (IBD) segments within pedigrees using observed marker data. Traditional linkage analysis requires specification of a trait model and can be misleading if this model is incorrect. Trait-model-free linkage methods are robust to model misspecification and address the challenge of unobserved IBD sharing directly. The trait-model-free linkage method of Basu et al. is similar to the transmission-based association statistics we investigate in that it is based on a preferential transmission model (PTM). PTMs are motivated by the idea that certain founder chromosomes harbor risk alleles at the trait locus and that these chromosomes are more likely to be transmitted to affected relatives compared to unaffected ones. Only parents with one nonbeneficial and one beneficial allele are informative in these PTM approaches. By contrast, traditional parametric linkage analysis makes use of all transmissions in the pedigree, making it more powerful than the PTM approach if the trait model is correctly specified. It would be interesting to compare the transmission statistics we have investigated to statistics from trait-model-free and traditional linkage methods, but such a comparison is beyond the scope of the present investigation.

A possible direction for future research is to extend the transmission probability model by including individual covariates such as sex or external information from bioinformatic predictions about the RVs. For example, covariates and disease subtype could be included in a logistic regression model for the transmission probability. However, the global or local configuration from one RV is unlikely to contain enough information to estimate any covariate effects in addition to the subtype-specific genetic effects. One approach to deal with the paucity of data from a single RV is to introduce random effects to borrow strength [Efron and Morris, 1972] across RVs. For example, a logistic regression model that includes *random* genetic effects and fixed covariate effects could be specified and fit to data from multiple RVs. Individual RVs can then be ranked by a ratio of posterior distributions of the random effects, in which the numerator is the posterior distribution evaluated at the posterior mean of the random effects, and the denominator is the posterior distribution evaluated at null values of the random effects [Fahrmeir et al., 2013]. One could include external biological information by allowing different random effects distributions for RVs in different classes; e.g., benign versus pathogenic as predicted by bioinformatics software such as PolyPhen2 [Adzhubei et al., 2010].

In conclusion, we propose new methods for prioritizing and testing rare variants in affected-only family designs, for diseases in which one subtype is more heritable than the others. To our knowledge, these methods are the first to incorporate information from disease subtypes. Our simulation results indicate that global likelihood approaches are robust to misspecification of the carrier probability for both prioritization and testing. For prioritization, likelihood methods perform better than RVS methods and are feasible for much larger studies. For testing, as expected, global likelihood methods generally have more power to detect *c*RVs than local methods, except when the carrier probability is grossly misspecified. The proposed methods will be useful in family-based sequencing studies seeking to prioritize rare variants for follow up with, e.g., pathway enrichment analyses or functional studies.

## 5 Author Contributions

CN and FBF developed the software and performed the analyses. CN, AB-W, AB and JG conceptualized the study. AB and JG provided supervison. CN, FBF and JG drafted the manuscript. All authors contributed to the editing and approval of the final manuscript.

## 6 Acknowledgments

CN and JG would like to pay their gratitude and respects to the late Janet Sinsheimer, our wonderful mentor, colleague and friend, without whose insights, support and encouragement this work would not have been possible. This research was funded by the Natural Sciences and Engineering Research Council of Canada (NSERC), the Canadian Statistical Sciences Institute (CANSSI) and the Canadian Institutes of Health Research (CIHR).

## 7 Conflict of Interest Statement

The authors declare no conflict of interest.

## 8 Data Availability Statement

The data and code that support the findings of this study are openly available from Zenodo at https://zenodo.org/records/10012025 [Nieuwoudt et al., 2023] and GitHub at https://github.com/simrvprojects.

## A Prioritizing RVs from 55 families

For testing RVs, the sample space of the reference distribution under the null hypothesis consists of all possible RV configurations, and so grows exponentially in the number of sequenced affected individuals. Global methods have 2^m^ − 1 possible global configurations, where m is the number of sequenced affected individuals in the study. Local methods have 2^k^ − 1 possible local configurations, where k is the number of sequenced affected individuals in families that carry the variant. As a result, computation of p-values for testing and ranking with the RVS-based methods, and for testing with the likelihood-based methods, is prohibitive for studies with more than a handful of families. However, ranking with the likelihood-ratio (LR) statistics requires only the observed RV configurations. By adding families successively we found that computation of the ranks scales linearly with the number of affected individuals in the study (Figure A.1). Consequently, likelihood-based ranking can be performed with much larger studies than the three families considered in the example analysis and the simulation study.

This Appendix presents a ranking analysis to prioritize the RVs on chromosome 8, but with a study comprised of all 55 families used in the simulations of Section 2.4. Computation of ranks for these data took less than 12 minutes on a laptop computer (Macbook Air with Apple M1 processor and 8GB of memory).

The 55 families contain a total of 855 affected individuals. Their simulated chromosome 8 data contains 1327 RVs, including all 10 *c*RVs from our pool of candidate *c*RVs. Of the 1327 RVs, 1167 (88%) appear in only one family, 123 (9%) appear in two families, 17 (1%) appear in three families, seven (0.5%) appear in four families and 13 (1%) appear in five or more families. In terms of affected individuals, 834 of the 1327 RVs (63%) appear in only one affected study member, 301 (29%) appear in two, 106 (8%) appear in three, 39 (3%) appear in four and the remainder (3%) appear in five or more affected individuals.

**Figure A.1:**
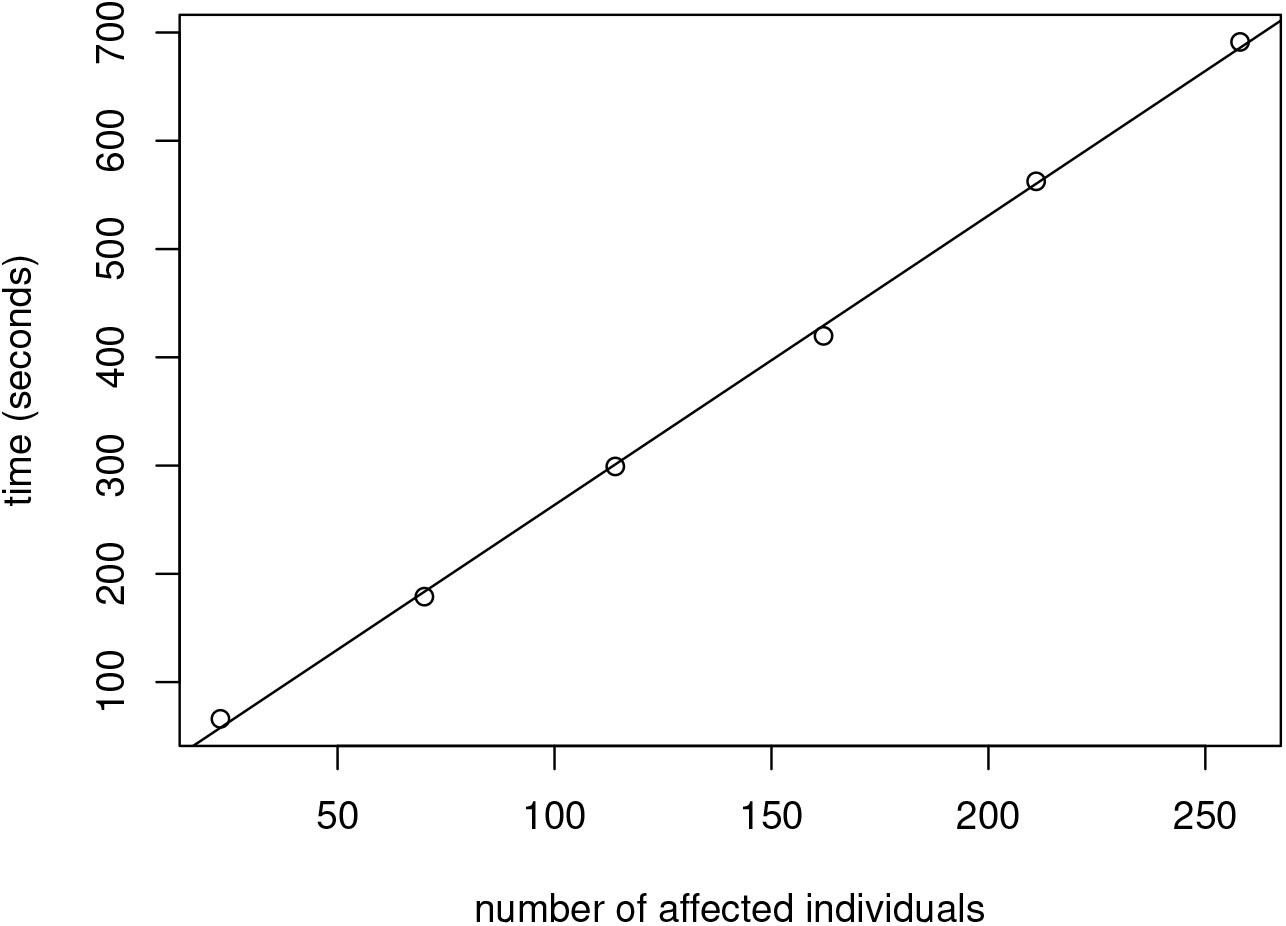
Time (in seconds) to compute ranks for all observed RVs *versus* number of affected individuals in the study. The different numbers of affected individuals were obtained by specifying studies of 5, 15, 25, 35, 45 or 55 families. Also shown is the least-squares regression line fit to these data.

The three log-likelihood ratio statistics were calculated for each observed configuration. For the two global statistics that require a value of the carrier probability, we used the true value of 0.00032 to simulate the data. For each LR statistic, we sorted their values and plotted against their rank to look for an “elbow” suggesting a break-point between a higher and lower priority group of RVs. Figure A.2 shows the plot for the top fifty local LR statistics. Plots for the two global LR statistics are similar. These plots suggest that the top-20 RVs might be considered higher priority.

**Figure A.2:**
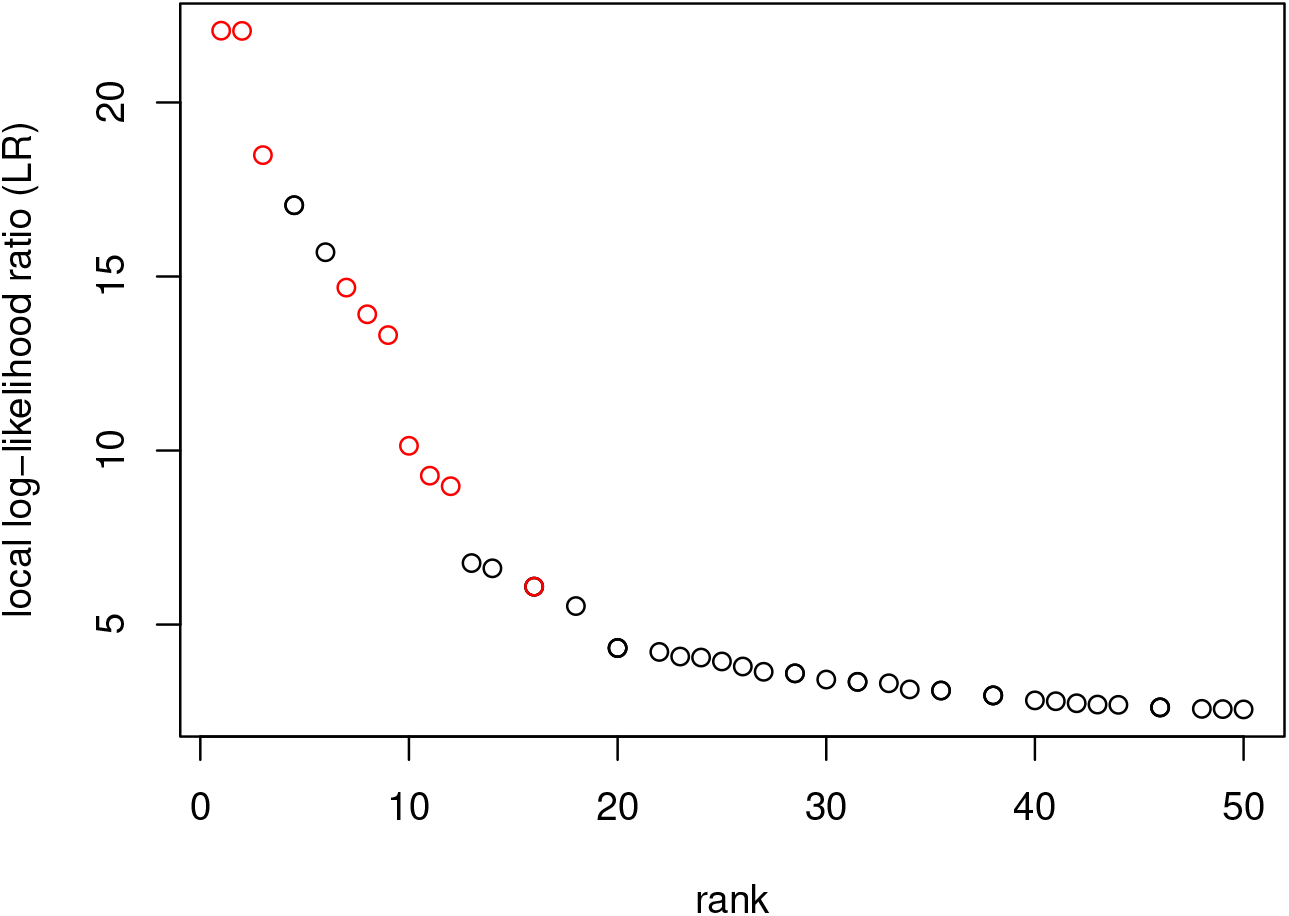
Local log-likelihood ratio (LR) statistics *versus* rank for the top-50 simulated RVs. Statistics for cRVs are colored red. Points for tied statistic values with the same average rank are overplotted.

A summary of the RVs with top-20 ranks is shown in Table A.2. All three LR methods give the same set of top-20 RVs. The local-LR ranks of the top-20 RVs are similar to those from the two global LR methods, except for three transposed pairs for RVs with ranks 1 and 2, 7 and 9, and 14 and 18. The correlations between the RV ranks for global and unconditional global LR, global and local LR, unconditional global and local LR statistics are, respectively, 0.88, 0.90 and 0.84. All 10 simulated cRVs are in the top-20 group, along with 11 non-causal RVs. The non-causal RVs appear in anywhere from one to eight families and, for all but five of these families, the non-causal RV is on the founder haplotype carrying the *c*RV, and tends to co-segregate with the *c*RV to affected individuals.

**Table A.2:** Top-20 ranked RVs, ordered by their chromosomal position, along with the number of segregating families (fams), number of affected carrier individuals (indiv) and rank from the three likelihood methods (global, unconditional global and local). The cRV column indicates whether the RV is simulated as causal (TRUE) or not (FALSE). The top-20 group of RVs is defined to be RVs with rank ≤20. RVs with tied values of the statistic are assigned the average rank.

| RV | cRV | RV count |  | rank |  |  |
| --- | --- | --- | --- | --- | --- | --- |
|  |  | fams | indiv | global | uncond | local |
| 8_7896589 | FALSE | 3 | 9 | 13.0 | 13.0 | 13.0 |
| 8_16140845 | FALSE | 3 | 8 | 16.0 | 16.0 | 16.0 |
| 8_17001992 | FALSE | 3 | 8 | 16.0 | 16.0 | 16.0 |
| 8_23020319 | TRUE | 2 | 8 | 11.0 | 11.0 | 11.0 |
| 8_23020346 | TRUE | 3 | 12 | 12.0 | 12.0 | 12.0 |
| 8_23020454 | TRUE | 8 | 30 | 7.0 | 7.0 | 9.0 |
| 8_23021159 | TRUE | 6 | 18 | 8.0 | 8.0 | 8.0 |
| 8_23021957 | TRUE | 3 | 8 | 16.0 | 16.0 | 16.0 |
| 8_23022671 | TRUE | 6 | 18 | 9.0 | 9.0 | 7.0 |
| 8_23068278 | TRUE | 9 | 33 | 1.0 | 1.0 | 2.0 |
| 8_23068413 | TRUE | 7 | 25 | 2.0 | 2.0 | 1.0 |
| 8_27433410 | FALSE | 8 | 28 | 4.5 | 4.5 | 4.5 |
| 8_27659221 | FALSE | 8 | 28 | 4.5 | 4.5 | 4.5 |
| 8_28494273 | FALSE | 8 | 26 | 6.0 | 6.0 | 6.0 |
| 8_39680176 | FALSE | 5 | 16 | 14.0 | 14.0 | 18.0 |
| 8_96609754 | FALSE | 4 | 11 | 20.0 | 20.0 | 20.0 |
| 8_97729036 | FALSE | 4 | 11 | 20.0 | 20.0 | 20.0 |
| 8_97994585 | FALSE | 4 | 11 | 20.0 | 20.0 | 20.0 |
| 8_118923860 | TRUE | 5 | 16 | 10.0 | 10.0 | 10.0 |
| 8_118933213 | TRUE | 6 | 22 | 3.0 | 3.0 | 3.0 |
| 8_143299677 | FALSE | 1 | 5 | 18.0 | 18.0 | 14.0 |

## Notes

https://doi.org/10.5281/zenodo.10012025

### Competing Interest Statement

The authors have declared no competing interest.

### Summary of Updates

Changes in response to reviewers' comments.

